# Rapid ‘multi-directed’ cholinergic transmission at central synapses

**DOI:** 10.1101/2020.04.18.048330

**Authors:** Santhosh Sethuramanujam, Akihiro Matsumoto, J. Michael McIntosh, Miao Jing, Yulong Li, David Berson, Keisuke Yonehara, Gautam B. Awatramani

## Abstract

Acetylcholine (ACh) is a key neurotransmitter that plays diverse roles in many parts of the central nervous system, including the retina. However, assessing the precise spatiotemporal dynamics of ACh is technically challenging and whether ACh transmits signals via rapid, point-to-point synaptic mechanisms, or broader-scale ‘non-synaptic’ mechanisms has been difficult to ascertain. Here, we examined the properties of cholinergic transmission at individual contacts made between direction-selective starburst amacrine cells and downstream ganglion cells in the retina. Using a combination of electrophysiology, serial block-face electron microscopy, and two-photon ACh imaging, we demonstrate that ACh signaling bears the hallmarks of both non-synaptic and synaptic forms of transmission. ACh co-activates nicotinic ACh receptors located on the intersecting dendrites of pairs of ganglion cells, with equal efficiency (non-synaptic)— and yet retains the ability to generate rapid ‘miniature’ currents (∼1 ms rise times: synaptic). Fast cholinergic signals do not appear to depend on anatomically well-defined synaptic structures. We estimate that ACh spread is limited to ∼1-2 *µ*m from its sites of release, which may help starbursts drive local direction-selective cholinergic responses in ganglion cell dendrites. Together, our results establish the functional architecture for cholinergic signaling at a central synapse and propose a novel motif whereby single presynaptic sites can co-transmit information to multiple neurons on a millisecond timescale.

## Main

Despite being intensely investigated over many decades, the functional and structural basis for cholinergic transmission in the central nervous system (CNS) remains obscure, partly because the methods required to precisely assess ACh dynamics in identified networks have not been available^1-6^. One body of literature supports the idea that conventional synaptic mechanisms mediate acetylcholine (ACh) signals, similar to GABAergic/glutamatergic transmission^7-12^. In this mode, ACh released from presynaptic nerve terminals is ‘directed’ to postsynaptic receptors positioned immediately opposite to the points of vesicle release (<0.3 *µ*m), enabling the generation of rapid responses^7,13-16^. On the other hand, competing evidence suggests that non-synaptic mechanisms (such as ‘volume’ transmission) mediate ACh signals^5,17-20^. Here, ACh diffuses over significant distances (> 1 *µ*m^14^) activating high-affinity receptors located at neighboring synapses with little spatial selectivity and temporal precision, much like classical neuromodulators (e.g., enkephalin, dopamine, and norepinephrine).

Electron microscopy studies have not provided a consensus regarding the extent to which cholinergic terminals form well-defined synaptic structures^1,2,18,21,22^. What complicates these results is the recent realization that many cholinergic neurons also release a second fast neurotransmitter (GABA^22-25^ or glutamate^19,26^), often from the same terminal varicosities^19,22,27^, making it difficult to determine the ultra-structural elements that are associated with cholinergic transmission. Results from functional studies also offer a limited view, as they lack the spatiotemporal resolution required to detect synaptic ACh signals (reviewed by^1,2^). Even ‘phasic’ postsynaptic cholinergic responses revealed in electrophysiological studies are relatively slow compared to transmission at conventional glutamatergic/GABAergic synapses^7,22,23,25,28^ (Fig. 1b; Extended Data Fig 1); and rarely associated with spontaneous ‘miniature’ events^9,12^, the standard defining feature of a chemical synapse^29^.

**Fig. 1:**
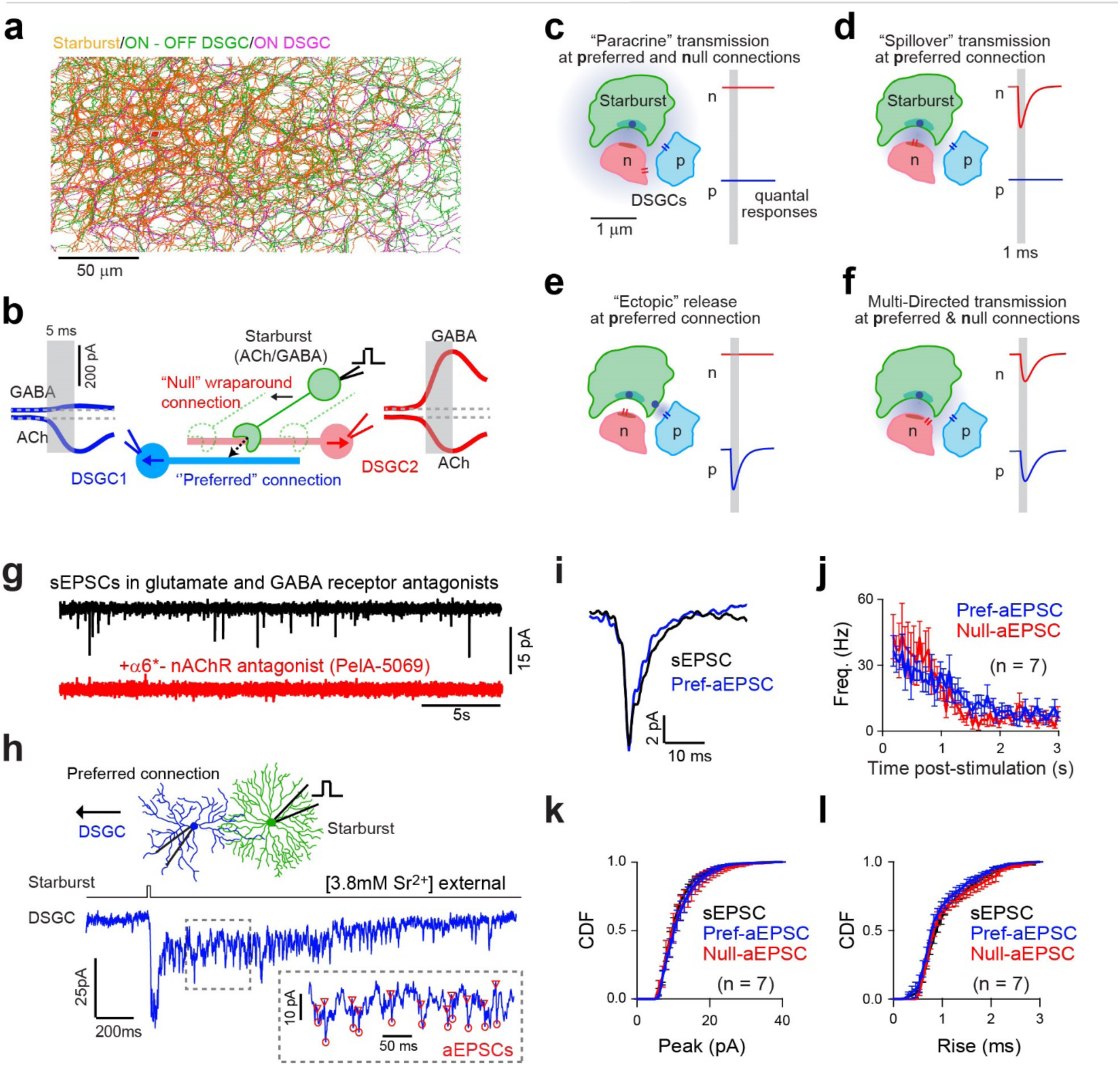
Quantal transmission between starbursts and DSGCs. **a.** An SBEM partial reconstruction of the ‘honeycomb’ cholinergic plexus formed by overlapping dendrites of densely spaced GABAergic/cholinergic starburst amacrine cells. Starburst release GABA and ACh varicosities that are distributed over their distal dendrites (not shown). **b.** Starburst dendrites respond preferentially to centrifugal motion (soma-to dendrite); their varicosities ‘wraparound’ and make strong synaptic contacts selectively with DSGCs encoding the opposite direction (DSGC2; arrows indicate directional preferences). These are considered ‘null connections’ because stimulation of the starburst through a patch electrode evokes large GABAergic response that dwarf the cholinergic responses co-activated in the synaptically connected DSGCs (DSGC2). Stimulation of the same starburst also evokes cholinergic (but not GABAergic) currents in DSGC encoding the same direction (DSGC1) through ‘preferred’ connections. The differential transmission of ACh/GABA aids in shaping direction selectivity^28,32-35^, but its cellular basis remains unknown. **c-f.** Putative models for cholinergic transmission, based on well-established synaptic/non-synaptic mechanisms^15,36-4215,36-4215,36-4215,36-4215,36-4215,36-4215,36-4215,36-4215,36-4215,36-4215,36-4215,36-4215,36-4215,36-4215,36-4215,36-4215,36-4215,36-4215,36-4215,36-4215,36-4215,36-4215,36-4215,36-4215,36-4215,36-4215,36-4215,36-4215,36-42^ predict different quantal responses at preferred and null-connections. **c.** ‘Paracrine’ transmission model^5,17-20^: ACh released from a single vesicle does not produce quantal responses. **d.** ‘Spillover’ transmission model^36,37,39,48^: ACh rapidly activates nAChRs only at null connections; some fraction also diffuses to neighbouring synapses to mediate slower, weaker responses at preferred connections. **e**. ‘Ectopic’ transmission model^36,37^: vesicle release occurs at sites that are not associated with the presynaptic specialization in the starburst varicosity (dark green), producing rapid responses at preferred connections, independent of release occurring at the null connections. **f**. ‘Multi-directed’ transmission model: ACh release from a single vesicle diffuses to both preferred and null sites. But unlike paracrine transmission, nAChRs can still be rapidly gated despite experiencing lower concentrations of the transmitter. **g.** Spontaneous cholinergic excitatory postsynaptic currents (sEPSCs) in a DSGC voltage-clamped at -60mV (E_Cl_), measured in control (top) and in the added presence of α6*-specific nAChR peptide (PelA-5069). **h.** Replacing extracellular Ca^2+^ with Sr^2+^ results in the desynchronization of vesicle release, providing insights into the unitary properties of preferred connections. Inset shows miniature-like fast asynchronous events (aEPSCs) observed after brief depolarization of the starburst to activate preferred connections. **i.** The waveforms of the average aEPSCs measured in h, overlaid with the average sEPSCs in the same cell. **j.** The instantaneous frequencies of responses mediated by preferred or null connections (n=7 pairs each). Data represented as mean ± SEM. **k, l.** Cumulative frequency distribution of the peak (k) and rise times (l) of the sEPSCs compared to the aEPSCs, elicited by preferred- and null-connections (n=7 each). Data represented as mean ± SEM.

In the retina, the primary sources of ACh are the direction-selective dendrites of cholinergic/GABAergic starburst amacrine cells (starbursts; Fig. 1a). Starburst dendrites stratify within two narrow bands in the inner retina^21^ and rely on volume transmission to convey cholinergic signals to neurons with processes in other layers of the inner retina^30,31^. Diffusion over long distances renders cholinergic signals non-directional, as ACh is accumulated from overlapping dendrites encoding different directions. However, within the cholinergic plexus, starburst dendrites also make large ‘wraparound’ synaptic contacts with specific direction-selective ganglion cells^21^ (DSGCs; Fig. 1b), through which they transmit rapid GABAergic and cholinergic signals^28,32-35^ (Fig. 1b). Wraparound synapses are ‘null connections’ that predominantly arise from starburst somas located on the ‘null’ side of the DSGC’s receptive field (i.e., the side from which null stimuli enter). Curiously, starbursts located on the DSGC’s preferred-side make significantly fewer synaptic contacts (<10%, and provide few GABAergic inputs) but still mediate robust cholinergic responses^28,32-35^ via ‘preferred connections’ (Fig. 1b).

The precise synaptic or non-synaptic mode (Fig. 1c-e)^15,36-42^ that the direction-selective circuit uses to transmit ACh signals remains unknown. Here we assess the spatiotemporal profile of ACh at single connections and show how it impacts the functional symmetry of the circuit to ultimately help shape direction-selectivity in DSGCs, which is currently a matter of debate^21,28,33-35,43-45.^

### Quantal ACh signals in the absence of anatomically well-defined synapses

In the most parsimonious model, ACh signals are mediated by volume transmission^21,28^, and insensitive to the specific wiring patterns (Fig. 1c). Contrary to this notion, however, we observed robust levels of spontaneous miniature excitatory postsynaptic currents EPSCs (sEPSCs) in voltage-clamped DSGCs (measured at a holding potential of -60 mV, in the presence of a drug cocktail containing antagonists for glutamate, and GABA receptors; Fig. 1g), likely reflecting single quantal events. Spontaneous EPSCs had an average peak amplitude of 10±2 pA, rapid kinetics (20–80% rise times: 1.2±0.1 ms; decay constant: 4.8±0.2 ms; n=10; Fig. 1i,k,l); and were almost entirely blocked by PelA-5069 (n=5), a peptide that specifically blocks α6 subunit-containing nicotinic ACh receptors^46^ (AChRs) (Fig. 1g). This indicated a predominant expression of α6 AChRs in DSGCs, consistent with single-cell transcriptomic profiling results^47^. The overt presence of miniature events suggests that at least part of the cholinergic signal is mediated by conventional point-to-point forms of synaptic transmission, and forced us to consider other possible synaptic configurations (Fig. 1d-f).

When compared to sEPSCs, the responses evoked by directly depolarizing starbursts through a patch electrode are relatively slow^28,32-34^ (rise times ∼5-10 ms; Extended Data Fig 1; Extended Table 1) and do not indicate the origin of sEPSCs. One hypothesis is that *bona fide* synaptic contacts (i.e., the ‘null’ connections) are the source of sEPSCs (Fig. 1b)^21^; and ACh transmission at preferred connections occur through ‘spillover mechanisms’, which would thus be weaker relative to null connections^33^, with quantal events likely falling below detection limits^36,37,39,48^ (Fig 1d). To study the properties of unitary connections, we desynchronized vesicular release from starbursts by replacing extracellular Ca^2+^ with Sr^2+^ (Fig. 1h)^49^. This led to the emergence of long-lasting barrages of asynchronous EPSCs (aEPSCs) that were similar to sEPSCs (Fig. 1h, i, k, l; Extended Data Table 1). Notably, the amplitudes, frequency, and kinetics of the isolated aEPSCs evoked via preferred and null connections (i.e., by depolarizing preferred and null-side starburst, respectively) were indistinguishable (Fig. 1j-l; Extended Data Table 1). These results indicate that the millisecond scale cholinergic signaling does not require wraparound synaptic contacts.

### Rapid ‘multi-directed’ transmission

In theory, the small number of ‘miswired’ starburst connections^21^, or vesicle release from ‘ectopic’ sites within the wraparound (i.e., release sites that function independently of the main presynaptic specialization^36,37^; Fig. 1e), could serve as the substrate for rapid cholinergic signals. The original SBEM datasets^21^ lacked ultrastructural details (vesicles) and would have overlooked putative ectopic sites. However, this did not appear to be the case because quantal inputs appeared to be shared. Simultaneous voltage-clamp recordings from pairs of DSGCs with overlapping dendritic fields (inter-somatic distance < 50μm; Fig. 2a) revealed that cholinergic sEPSCs occurred synchronously (12 ± 1%; n=7 pairs; Fig. 2b, c; Extended Data 2a-c), more frequently than expected from their spontaneous rates (∼3Hz). Correlated sEPSCs directly demonstrated that ACh released from single vesicles can diffuse to multiple sites and yet rapidly activate synaptic-like currents, providing direct evidence for a novel mode of ‘multi-directed’ transmission (Fig. 1f). This resembles volume transmission with one major difference—signals occur on a millisecond timescale.

**Fig. 2:**
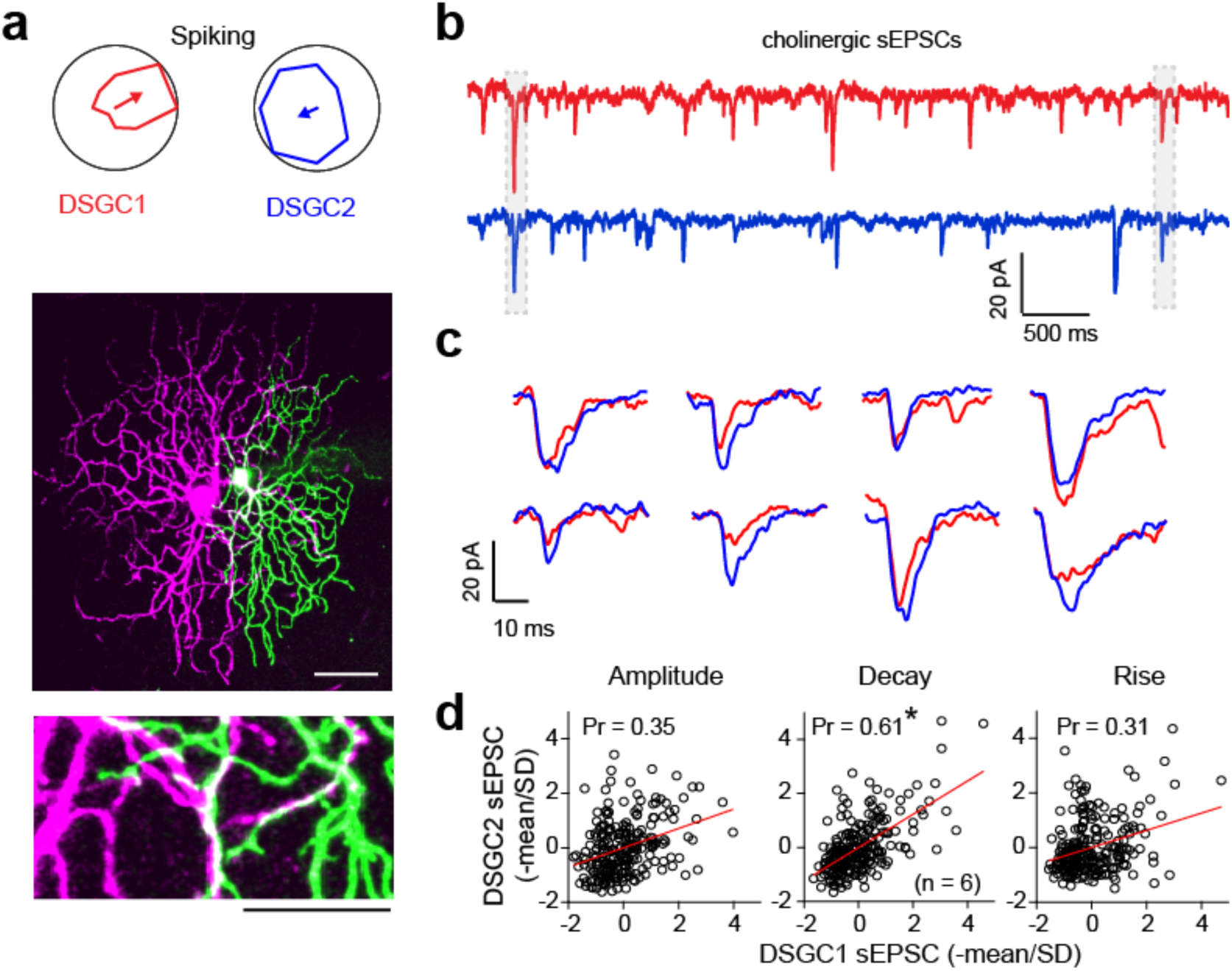
Rapid, multi-directed cholinergic transmission. **a.** Polar plots depicting the magnitude of the spiking responses recorded from a pair of adjacent DSGCs (top). The dendritic morphologies of the same pair of DSGCs (*middle*; scale bar 50μm). A magnified view of a small section of the image where the dendrites of the two DSGCs co-fasciculate (*bottom*; scale bar 25μm). **b.** Cholinergic sEPSCs recorded simultaneously in the two nearby DSGCs demonstrate that a significant fraction of sEPSCs are synchronized (boxed regions). Thus, ACh release from single vesicles can rapidly activate two sites i.e., ACh is multi-directed. **c.** Sample of synchronous cholinergic sEPSCs shown on an expanded timescale. **d.** The peak amplitudes (*left*), decay constants (*middle*) and rise times (*right*) of the individual events that occurred synchronously are plotted against each other (241 events from 6 pairs). Points indicate the deviation of the event parameter from the mean (normalized to the standard deviation). (*p < 0.001, see methods).

Several control experiments and analyses bolster these conclusions. First, GABAergic inputs in pairs of DSGCs were not correlated, confirming that synapse specificity is maintained at inhibitory synapses (n=4; Extended Data 2d-f), which served as a positive control for our methodology. Second, sEPSCs and sIPSCs were not correlated, arguing that although GABA and ACh vesicles may be present in the same varicosities, they are released independently. Thus, the release of multiple vesicles from single starburst varicosities (or from more than one varicosity on a given starburst dendrite) is not likely to occur under resting conditions (Extended Data Fig. 2g-i). Third, distribution of the peak amplitude of the correlated sEPSCs measured in a DSGC was similar to the population distribution of sEPSCs, indicating that correlated events did not represent a select population of larger events associated with multivesicular release^50^ (Extended Data Fig. 3; Extended Data Table 1). Thus, it is likely that ACh released from single starburst varicosities spread to co-activate nAChRs at two postsynaptic sites.

Nicotinic AChRs mediating synchronous sEPSCs are likely to be located at some distance away from the release site, as they share a common source of ACh. They are likely to experience slower, lower concentration ACh transients compared to receptors positioned directly across the release site. Close inspection of individual pairs of correlated sEPSCs recorded in adjacent DSGCs revealed a striking correlation in their decay kinetics (Fig. 2c,d), and to a lesser extent, in their rise-times and amplitudes. This raised the possibility that local ACh concentration profiles, rather than the unbinding of the transmitter from receptors (i.e., deactivation) shape the decay kinetics of cholinergic sEPSCs^15^. Consistent with this notion, blocking ACh esterase (50 nM ambenonium), which slows the clearance of ACh by preventing its hydrolysis, prolonged the decay phase of sEPSCs (Extended Fig. 4). In this context, nAChRs behave similarly to ‘extrasynaptic’ NMDA receptors whose activity is shaped by low micromolar levels of glutamate experienced during spillover transmission and enhanced when glutamate clearance is compromised^40-42,51^—except nAChRs have significantly faster activation kinetics.

**Fig. 4:**
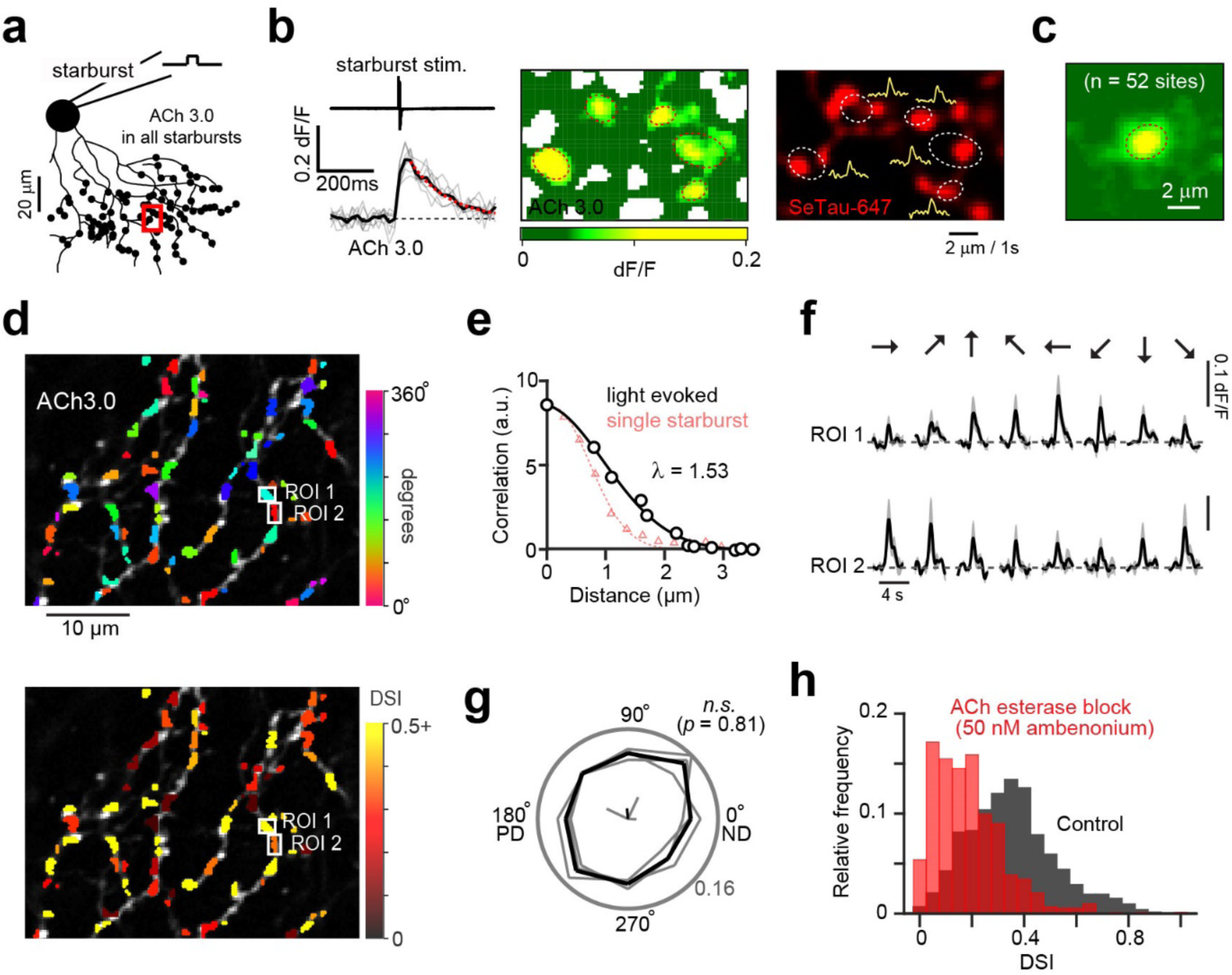
Two-photon imaging of ACh using a genetically encoded senor reveals that ACh is locally, but not globally tuned for direction. **a.** A genetically encoded ACh sensor (ACh3.0) was selectively expressed in many starbursts (see Extended Data Fig. 6), one of which was stimulated with a patch electrode. A red indicator (SeTau-647) was included in the electrode solution, to reveal its morphology. **b.** Depolarizing the starburst produced transient excursions in ACh3.0 fluorescence (*left*; red dotted line represents an exponential fit to the response decay) that were spatially localized (*middle*; the dotted line indicates the 1 standard deviation level contour of a 2D Gaussian fit) near starburst varicosities imaged in the red channel (*right*; yellow traces indicate the filtered response at each site). **c.** The average spatial profile for 52 sites measured in 8 starbursts. **d.** Two photon image of DSGC dendrites selectively expressing ACh3.0, in the Oxtr-T2A-Cre mouse line (Extended Data Fig. 8). The colors indicate the preferred direction for each site (*top*) and its tuning strength quantified using the direction selectivity index (DSI; 0 indicates non-tuned, 1 indicates perfect tuning; *bottom*). **e.** Noise correlations (trial-to-trial fluctuations around the mean ACh3.0 signal) plotted over distance. The spatial decay of ACh signals associated with single starburst stimulation measured in **c** is shown for comparison (red triangles). **f.** Example ACh signals evoked by spots moving in 8 different directions, measured in the two ROIs shown in **d**. Data represented as mean ± SEM from 4 trials. **g.** Relative frequency histogram of the preferred directions of all the ROIs in polar form (*p*=0.81, n=723 ROIS, 3 DSGCs; Hodges-Ajne non-parametric test for angular uniformity of the data) indicates a random distribution of angles with respect to the DSGCs preferred and null directions (PD and ND, respectively). **h.** Relative frequency histogram of the DSIs of all the ROIs in control, and in the presence of an ACh esterase blocker (50 nM ambenonium).

To estimate how far ACh spreads from its point of release, we compared the degree to which sEPSCs were synchronized to the level of dendritic overlap between the recorded pairs of DSGCs^52^. Anatomical analysis of the dendritic morphologies (recovered after the physiological recordings) showed that the fraction of dendrites of a given DSGC, that were within a certain distance of the dendrites of a second DSGC, grew rapidly as this distance increased (‘nearest neighbor analysis’; See methods; Fig. 3a, b). If ACh spread across a radius of 5*µ*m, it would have a∼50% chance of coactivating receptors on two dendrites simultaneously. However, the fraction of correlated events measured experimentally was only 12±1%, which predicts ACh spreads ∼1*µ*m from its site of release (Fig. 3c). It is important to note that this analysis only provides an average estimate of the spread and hinges on the assumption that many starburst wraparound synapses are proximal to the dendrites of a DSGC encoding a different direction, an assumption which we examined next using SBEM.

**Fig. 3:**
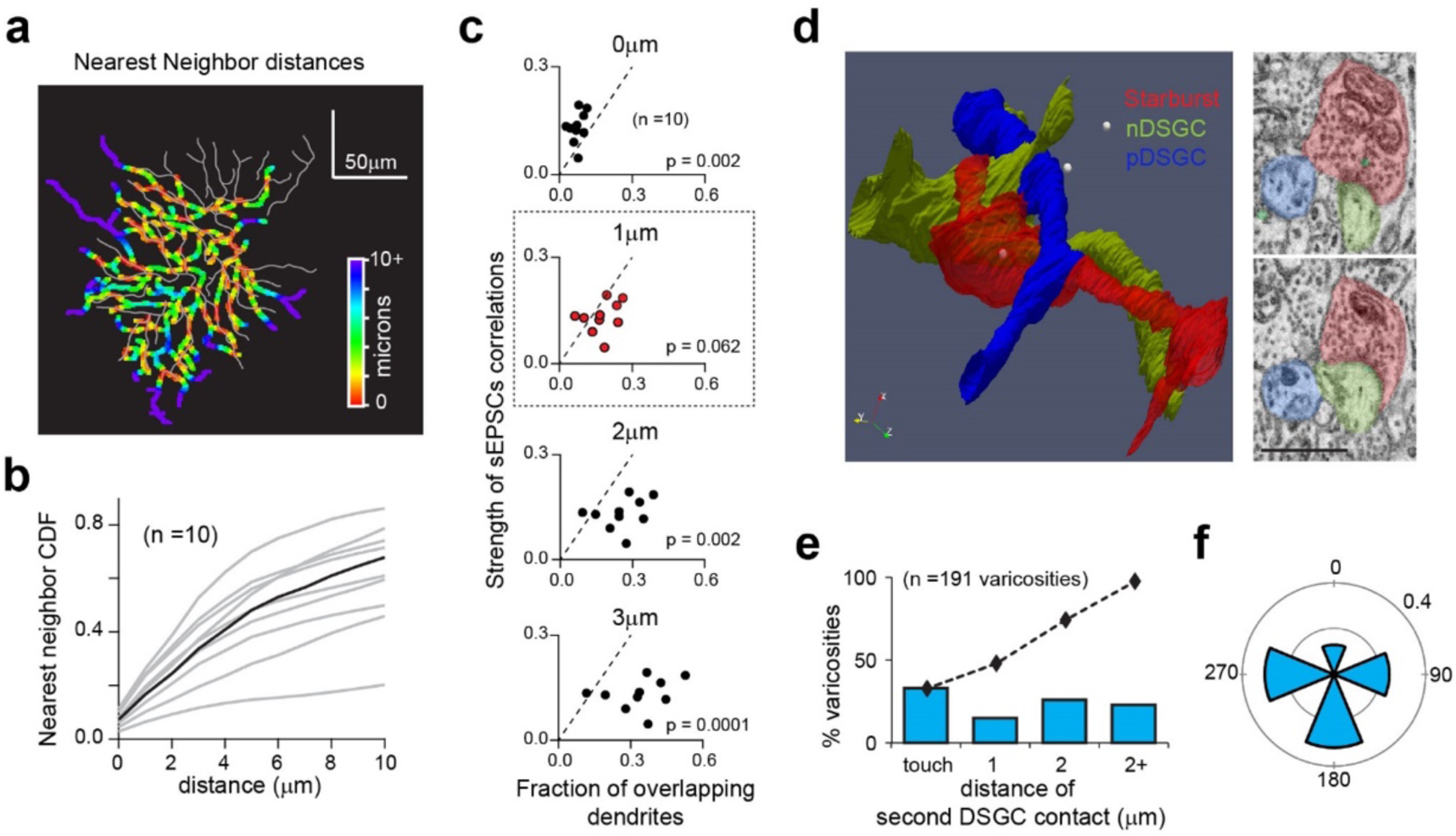
Multi-directed transmission is associated with local ACh signals, mediated by ‘tripartite’ complexes. **a.** Map of a DSGC’s dendritic arbor color coded to represent the distance of the nearest dendrite of a neighbouring DSGC (gray). **b.** As DSGCs co-fasciculate, the cumulative distribution function (CDF) of nearest neighbour distances grows rapidly with distance over which dendrites are considered to be neighbours (n=10 DSGCs from 5 pairs). Black trace indicates the CDF for cell shown in a. **c.** Comparison of the correlation strength of the sEPSCs recorded in DSGC pairs (Extended Data Fig. 2a-c) to the fraction of their dendritic overlap, for four different distances (0, 1, 2 and 3 “m; dotted line is the unity line). Correlations are well predicted when sites within 1 “m are considered neighbours i.e. share ACh released by the same vesicle. **d.** 3D SBEM reconstructions illustrating a ‘tripartite’ complex comprising a starburst varicosity and two DSGC dendrites. The starburst varicosity makes a wraparound null contact (white spheres) with one DSGC (nDSGC), and a peripheral contact with a second DSGC encoding the opposite direction (pDSGC) (see Extended Data Fig. 5). Panels on the right show cross-sections, illustrating the ultrastructure of the tripartite complex. Scale bar=1.5μm. **e.** Distribution of distances between a starburst varicosity and its closest peripheral contact. Dotted line shows the CDF. **f.** The directional preference of the DSGCs receiving peripheral contacts (n=125 varicosities), relative to the DSGC receiving the wraparound contacts.

In an SBEM stack^53^ (50 × 210 × 260*µ*m^3^), we reconstructed 82 ON starburst cells in sufficient detail to infer the directional preferences of their dendrites based on the soma location (Fig. 1a; Extended Data Fig. 5). We also reconstructed dendrites arising from 32 ON-OFF DSGCs, of which only half had somas in the volume. By mapping the asymmetry of ON starburst synaptic contacts onto the DSGCs, we could divide the DSGCs into four groups, each corresponding to the encoding of single cardinal direction^21^ (Extended Data Fig. 5). To assess the abundance of other DSGC dendrites in the neighborhood of starburst-to-DSGC synapses, we began by mapping a small fraction of classic starburst ‘wraparound’ synapses (n=191) onto six DSGCs with different directional preferences. Then, we measured the distance from each of these synapses to the closest point on the surface of any other ON-OFF DSGC. In nearly a third of the cases, the varicosity was in direct contact with the dendrite of another DSGC (Fig. 3d). In 77% of the total cases, there was a dendrite of another DSGC within 2 *µ*m (Fig. 3e). The directional preference of the nearest-neighbor DSGC bore no consistent relationship to that of the DSGC receiving the wraparound synapse (except for a tendency to avoid secondary connections with neighboring DSGCs encoding the same direction; Fig. 3f).

We also made an exhaustive reconstruction of the SAC-DSGC network in a smaller volume (7 × 7 × 7 *µ*m^3^; Extended Data Fig. 6), comprised of processes arising from 5 ON-OFF DSGCs, 2 ON DSGCs and 43 ON starbursts. Starburst dendrites encoding all possible directions were represented in the volume, with no apparent spatial order (Extended Data Fig. 6a). To assess the proximity of DSGC dendrites to starburst release sites, we measured the distance between each of many points on the surface of DSGC dendrites in the middle of the volume and the nearest starburst wraparound synapse onto any other cell (regardless of cell type). The mean distance was 0.8 *µ*m (Extended Data Fig. 6b). Because the reconstruction was centered on a dense aggregation of ON starburst processes, this number should be taken as a lower bound for the separation of starburst varicosities from their non-wraparound targets.

As with many cholinergic synapses elsewhere in the CNS^17,18,22^, putative secondary peripheral contacts made by starburst wraparounds were devoid of pre- or postsynaptic specializations, making them difficult to verify by anatomy alone (Fig 3d). It is only in the context of our electrophysiological data showing rapid synchronous cholinergic sEPSCs that we can show that transmission likely occurs at such ‘tripartite complexes’ (i.e., varicosity and two DSGC dendrites; Fig 3d).

### ACh is locally, but not globally, tuned for direction

Electrophysiological analysis indicates that under different conditions, cholinergic signals mediated by preferred connections appear to be stronger^32,45^, weaker^33^, or equal in magnitude^35,43,44^ (Fig. 1k) relative to those mediated by null connections. However, these results are generally difficult to interpret due to space-clamp considerations^54^; and as such, do not provide an indication of the patterns of excitation on single dendrites of the DSGC, where direction is computed ^28,55-57^. To circumvent these issues, we assessed ACh dynamics using a G-protein-coupled receptor activation-based sensor (ACh 3.0), which has an ACh binding affinity of ∼2 *µ*M, that is roughly comparable to that of the endogenous α6* nAChRs^58,59^.

To determine the spatial extent of ACh released from single starburst varicosities, we expressed ACh3.0 selectively in the starburst honeycomb (see Methods) and loaded single starbursts expressing the sensor with SeTau-647 (red indicator) through a patch electrode. Brief depolarizations of single starbursts evoked spatially localized, fluorescence transients (0.78±0.03 *µ*m; *τ*_decay_=130±13 ms; n=52 sites; Fig. 4a-c) that mapped to individual varicosities of the stimulated starburst, but were not necessarily centered on them (Fig. 4b; Extended Data Fig. 7). Responses could only be detected in ∼20% of the imaged varicosities (n=253 varicosities from 8 starbursts), suggesting non-uniform release properties of ACh across starburst dendrites.

During natural stimulation of the circuit with moving light bars, however, the contribution of individual varicosities to the ACh response was less evident, regardless of whether we expressed ACh3.0 selectively in starburst or DSGC dendrites (using the Chat-Cre and Oxtr-T2A-Cre mouse line that label starburst and nasally-tuned ON-OFF DSGCs, respectively; Extended Data Fig. 6 & 7). Nevertheless, an examination of the trial-to-trial variability of ACh3.0 responses revealed noise correlations (i.e., the variability around the mean response) that decayed over short distances (λ∼1.5 *µ*m; Fig. 4e; see Methods), likely reflecting a dominant contribution of ACh released from single starburst varicosities (Fig. 4e).

Signals measured in small regions of interest (ROIs, over which noise was correlated; see Methods) were strongly tuned for different directions (Fig. 4d, f, h; direction selectivity index [DSI]=0.35±0.01; n= 723 ROIs; See methods), indicating ACh release from varicosities was directional, similar to their calcium responses^60,61^. Blocking ACh esterase using 50 nM AMB, however, dramatically reduced the direction tuning at most sites (Fig. 4h; DSI=0.19±0.01; *p*=3.03×10^−65^, Wilcoxon signed-rank test). Thus, ACh esterase limits the spread of ACh to within ∼1-2 *µ*m of the release site and helps maintain the synaptic specificity of starburst cholinergic input during natural patterns of activity.

Color maps of the DSI and preferred angles show that the strength of the direction tuning was heterogeneous across the dendritic tree (Fig. 4d), and had no clear relation with the DSGC’s preferred-null axis (Fig. 4g), the latter indicated by its spiking response (Extended Data Fig. 8c). Besides, the average peak amplitudes of the ACh signals across the dendritic tree were non-directional (Extended Data Fig. 8g). It is important to note that the organization of the presynaptic wiring reflected in the imaging experiments is not contingent on the precise spatiotemporal profiles of ACh we measured, which may be distorted to some extent by the slow kinetics of the ACh sensor (Fig. 4b). The lack of global order in the tuning distribution or strength of local ACh3.0 signals, argues against the idea that cholinergic excitation is biased towards the DSGC’s preferred direction^32,45^.

At first glance, the lack of global order in the cholinergic excitation appears consistent with existing models in which direction selectivity arises from the integration of non-directional excitation and directionally tuned inhibition^21,34,35,44,55-57,61^. However, several lines of evidence suggest that non-linear integration of synaptic inputs occurs over subregions of the DSGC’s dendrite^28,57^, which can be as small as ∼10 *µ*m in length^55^. In this case, directional cholinergic input would shape local dendritic responses. When cholinergic inputs align with the null direction (i.e., the wraparound connections), the co-transmission of GABA occurring at that same site would effectively cancel excitation. Fine-scale coordination of excitation and inhibition is conditional on ACh signals being highly compartmentalized and directional. For cholinergic inputs aligned with the preferred direction, the chances that they receive coincident GABA input is low. In this way, the net cholinergic drive emerges strongly directional. Locally tuned cholinergic input implies a new role for ACh in the direction-selective dendritic computation.

Our electrophysiological and imaging results demonstrate the most spatiotemporally compartmentalized ACh signals recorded in the CNS to date (∼1 ms rise times; λ∼1.5 *µ*m). Yet these cholinergic signals do not appear to be associated with electron-dense postsynaptic specializations, nor do they require tight coupling between pre- and postsynaptic elements. From an anatomical perspective, cholinergic transmission appears to be mediated by volume transmission. In contrast, from a physiological perspective, it bears the hallmark of conventional synaptic mechanisms, thus suggesting a fundamentally different framework, i.e., multi-directed cholinergic transmission. Under this revised framework, the debate regarding the nature of cholinergic transmission, which has engaged neuroscientists for decades, appears fundamentally flawed in its tacit assumption that cholinergic neurons utilize similar mechanisms as other neurotransmitters or neuromodulators.

Multi-directed transmission is a unique and efficient way in which small populations of cholinergic neurons can powerfully modulate network activity with temporal precision, through their sparsely distributed connections^7,9-12,62,63^. Indeed, ‘local’ paracrine mechanisms have been proposed in other regions of the brain^7,22,23^, but confirming whether transmission is multi-directed in these areas remains a challenge for future investigations. The conceptual and technical approaches presented here set the stage for testing this hypothesis.

## Methods

### Animals

Electrophysiological experiments were performed using adult Hb9-EGFP (RRID: MGI_109160) or Trhr-EGFP (RRID: MMRRC_030036-UCD) crossed with ChAT-IRES-Cre (RRID: MGI_5475195) crossed with Ai9 (RRID: MGI_3809523). Acetylcholine imaging experiments were performed using Oxtr-T2A-Cre (strain: Cg-*Oxtr*^*tm1.1(cre)Hze*^/J, Jackson laboratory stock: 031303) and ChAT-IRES-Cre mice under C57BL/6J background. All mice were between two and sixteen months old in either sex. All procedures were performed in accordance with the Canadian Council on Animal Care and approved by the University of Victoria’s Animal Care Committee or in accordance to Danish standard ethical guidelines and were approved by the Danish National Animal Experiment Committee (Permission No. 2015-15-0201-00541).

### Physiological Recordings

Mice were dark-adapted for approximately 30–60 min before being briefly anesthetized and decapitated. The retina was extracted and dissected in Ringer’s solution under infrared light. The isolated retina was then mounted on a 0.22 mm membrane filter (Millipore) with a pre-cut window to allow light to reach the retina, enabling the preparation to be viewed with infrared light using a Spot RT3 CCD camera (Diagnostic Instruments) attached to an upright Olympus BX51 WI fluorescent microscope outfitted with a 40× water-immersion lens (Olympus Canada). The isolated retina was then perfused with warmed Ringer’s solution (35–37 °C) containing 110 mM NaCl, 2.5 mM KCl, 1 mM CaCl_2_, 1. 6 mM MgCl_2_, 10 mM dextrose and 22 mM NaHCO_3_ that was bubbled with carbogen (95% O_2_:5% CO_2_). Perfusion rates were maintained at ∼3 ml/min.

DSGCs were identified by their genetic labeling or by their characteristic direction-selective responses. Spike recordings were made with the loose cell-attached patch-clamp technique using 5 -10 M$ electrodes filled with Ringer’s solution. Voltage-clamp whole-cell recordings were made using 4 -7 MΩ electrodes containing 112. 5 mM CH3CsO3S, 7. 75mM CsCl, 1 mM MgSO4, 10 mM EGTA, 10 mM HEPES and 5 mM QX-314-Cl. For stimulating neurotransmitter release, starbursts were voltage-clamped with electrodes containing 112. 5 mM CH3CsO3S, 7. 75 mM CsCl, 1 mM MgSO4, 0.1 mM EGTA, 10 mM HEPES, 10 mM AChCl, 4 mM ATP, 0.5 mM GTP, 1 mM ascorbate and 10 mM phosphocreatine. The pH was adjusted to 7.4 with CsOH. In some paired recordings, 100*µ*M Alexa-488 or 10*µ*M SeTau-647 (SETA Biochemicals, #K9-4150) was added to the internal to visualize the dendritic morphologies. The reversal potential for chloride was calculated to be –56 mV. The junction potential was calculated to be -8 mV and was not corrected. Recordings were made with a MultiClamp 700B amplifier (Molecular Devices). Signals were digitized at 10 kHz (PCI-6036E acquisition board, National 9 Instruments) and acquired using custom software written in LabVIEW. Unless otherwise noted, all reagents were purchased from Sigma-Aldrich Canada. Ambenonium chloride and UBP310 were purchased from ABCAM Biochemicals. DL-AP4, SR-95531 and CNQX were purchased from Tocris Bioscience.

### Virus injections

The plasmid pssAAV-2-hSyn1-chI-dlox-Igk_(rev)-dlox-WPRE-SV40p(A) was generated by Viral Vector Facility at University of Zurich based on pAAV-hSyn-G that has been deposited to Addgene (#121922)^58^. The single-stranded AAV vector ssAAV-9/2-hSyn1-chI-dlox-Igk_(rev)-dlox-WPRE-SV40p(A) (1.2 × 10^13^ vg/ml) was produced using the plasmid by Viral Vector Facility at University of Zurich. For intravitreal viral injections of the AAV, mice were anesthetized with an i.p. injection of fentanyl (0.05 mg/kg body weight; Actavi), midazolam (5.0 mg/kg body weight; Dormicum, Roche) and medetomidine (0.5 mg/kg body weight; Domitor, Orion) mixture dissolved in saline. We made a small hole at the border between the sclera and the cornea with a 30-gauge needle. Next, we loaded the AAV into a pulled borosilicate glass micropipette (30 *µ*m tip diameter), and 2 *µ*l was pressure-injected through the hole into the vitreous of the left eye using a Picospritzer III (Parker). Mice were returned to their home cage after anesthesia was antagonized by an i.p. injection of a flumazenil (0.5 mg/kg body weight; Anexate, Roche) and atipamezole (2.5 mg/kg body weight; Antisedan, Orion Pharma) mixture dissolved in saline and, after recovery, were placed on a heating pad for one hour.

### Two-photon ACh imaging

Three to four weeks after virus injection into the eyes of Oxtr-T2A-Cre or ChAT-Cre mice, we performed two-photon imaging of ACh3.0 fluorescence signals on dendrites of the ON-OFF DSGCs or starbursts. Retinas were prepared similar to the physiological recordings. The isolated retina was then placed under the microscope (SliceScope, Scientifica) equipped with a galvo-galvo scanning mirror system, a mode-locked Ti: Sapphire laser tuned to 940 nm (MaiTai DeepSee, Spectra-Physics), and an Olympus 60× (1.0 NA) objective. The ACh3.0 signals emitted were passed through a set of optical filters (ET525/50m, Chroma; lp GG495, Schott) and collected with a GaAsP detector. Images were acquired at 8-12 Hz using custom software developed by Zoltan Raics (SELS Software). The size of the imaging window was 40-60 *µ*m in length and 80-120 *µ*m in width (0.45 *µ*m/pixel). Temporal information about scan timings was recorded by TTL signals generated at the end of each scan, and the scan timing and visual stimulus timing were subsequently aligned during off-line analysis. In experiments where responses were evoked via depolarizing starbursts, ACh responses were measured at higher temporal resolution (∼50Hz)^55^, which enables a more accurate assessment of the decay kinetics of the response (Fig. 4b). In these experiments, the size of the imaging window was typically 30*µ*m × 20*µ*m (0.28 *µ*m/pixel; dwell time=1*µ*s/pixel).

### Imaging data analysis

For analyzing the direction-selective properties of the ACh3.0 signals, regions of interest (ROIs) for acetylcholine signals were determined by customized programs in MATLAB. First, the stack of acquired images was filtered with a Gaussian filter (3 × 3 pixels), and then each image was down-sampled to 0.8 of the original using a MATLAB imresize function. The signals in each pixel were smoothed temporally by a moving average filter with a window size of 2 time-bin and resampled using the MATLAB interp function. To evaluate the spatial scales of the light-evoked acetylcholine signals, we calculated the noise correlation among the signals in pixels during static flash (300 *µ*m diameter, 50% contrast), and plotted the correlation coefficient against the distance between pixels (Fig. 4e). The correlation was normalized as a score (C):

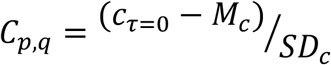

where *c*_*τ*=0_ is the amplitude at time delay τ=0 in cross-correlation function between pixel *p* and *q, M*_*c*_ and *SD*_*c*_ are mean and SD of the cross correlation function, respectively. The spatial scale of ACh responses were estimated by fitting the decay of noise correlations to a 2-D Gaussian function in MATLAB (Fig. 4e). We set a threshold of the correlation score, > 2.5, to determine which pixels were to be included as a single ROI. The response of each ROI (Δ*F*(*t*)) was calculated as

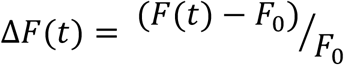

where *F*(*t*) is the fluorescent signal in arbitrary units, *F*_0_ is the baseline fluorescence measured as the average fluorescence in a 1 second window before the presentation of the stimulus. After the processing, responsive pixels were detected based on a response quality index (RI):

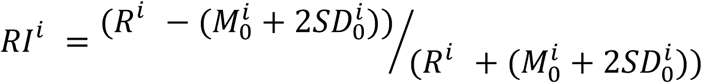

where *R*^*i*^ is a peak response amplitude during motion stimulus to direction *i*, 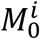 and 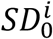 are mean and SD of ACh signals before stimulus (1 s period), respectively. The ROIs with the RI higher than 0.6 were determined as responsive (Extended Data Fig. 7e).

To evaluate the directional tuning, we calculated direction selectivity index (DSI):

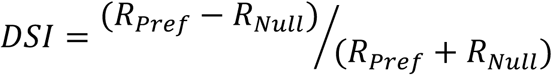

where *R*_*Pref*_ and *R*_*Null*_ denote the amplitude of ACh3.0 signals to preferred and null direction, respectively. DSI ranged from 0 to 1, with 0 indicating a perfectly symmetrical response, and 1 indicating a response only in the preferred direction. The preferred direction for individual ROIs was defined as an angle (*θ*) calculated by the vector sum:

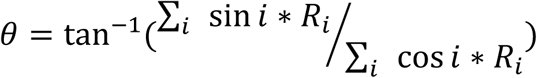

where *i* denotes the motion direction, and *R*_*i*_ denotes the response amplitude.

### Light stimuli

Visual stimulation was generated via custom-made software (Python and LabVIEW) by Zoltan Raics. The generated stimulus was projected using a DLP projector (LightCrafter Fiber E4500 MKII, EKB Technologies) coupled via a liquid light guide to an LED source (4-Wavelength High-Power LED Source, Thorlabs) with a 400 nm LED (LZ4-00UA00, LED Engin) through a band-pass optical filter (ET405/40×, Chroma). The stimulus was focused on the photoreceptor layer of the mounted retina through a condenser (WI-DICD, Olympus). The stimuli were exclusively presented during the fly-back period of the horizontal scanning mirror^56^. To measure the directional tuning, we used a spot (300 *µ*m diameter, 2 s duration) moving in 8 directions (at 800-1200 *µ*m/s) with a 50 % contrast.

### Anatomical Analysis

After estimating the sEPSC correlations, the dendritic morphology of the DSGCs was visualized with Alexa-488 and SeTau-647 using two-photon imaging techniques. The skeletons of the ON and OFF dendritic trees of the DSGCs were traced using a custom written script in IGORPro and were analyzed separately. The ON and OFF arbours were flattened to a single plane in which all dendrites in this analysis had a uniform dendritic thickness of 1 *µ*m. From these images, we estimated dendritic overlap as previously described^52^ (Fig. 3b).

For ultrastructure, we used a previously published SBEM^53^ data set (retina k0563). Voxel dimensions were 12 × 12 × 25 nanometer (nm) (x, y, and z, respectively). Potential DSGCs were first identified as ganglion cells with bistratified morphology in the IPL. Next, wrap-around synapses from starbursts on the dendrites of these cells were identified by their electron-dense membrane thickening laden with vesicles (Fig. 3d); reconstruction of the starbursts confirmed their co-stratification with the ganglion cell dendrites identifying it as a DSGC (Extended Data Fig. 4). Next, to estimate the closest peripheral contact, we retraced the dendrites of all the identified DSGCs in the volume (Fig. 1a), and examined the closest contact made by these DSGC dendrites to starburst varicosities outside the wrap-around synapse.

In order to estimate the average distance of DSGC dendrites from starburst release sites (varicosities), a 7×7×7*µ*m^3^ region was selected, where most of the ON starburst dendrites were reconstructed (Fig. 1a; Extended Data Fig. 5). Interestingly, we noted that along with the classical single varicosities, there were a few double varicosities (directed to two separate DSGC dendrites). Once, all the synapses were annotated, we measured the distance between each of many points on the surface of DSGC dendrites in the middle of the volume and the nearest classic wraparound synapse SAC onto any other cell, regardless of cell type. All analyses were performed by tracing skeletons and annotating synapses using the Knossos software package (www.knossostool.org). Volumetric reconstructions of synapses were performed using ITK-SNAP (www.itksnap.org).

### Spontaneous EPSC analysis

Asynchronous EPSCs was acquired from DSGCs for 1 second post-starburst stimulation, while spontaneous EPSCs in the same DSGC was acquired for 20 seconds after the aEPSCs. Preferred- and null-starburst-DSGC connections were identified by estimating the preferred direction of the DSGC using extracellular spike recordings before performing paired recordings. In cases where only sEPSCs were acquired, DSGCs were recorded in a dark background, for a period of 5-15 minutes. Once acquired, the traces were low pass filtered at 300 Hz, and fast-rising events were detected automatically using tarotools (sites.google.com/site/ tarotoolsregister/) in IgorPro. The events were detected using an amplitude thresholds set between 5 – 7 pA. Multiple overlapping events observed during the asynchronous activity were separated using the default (2.5ms) decay constant parameter^49^. Peak amplitude and 20-80% rise times were calculated with in-built tarotools functions.

### Statistical testing

Population data have been expressed as mean ± SEM and are indicated in the figure legend along with the number of samples. Student’s t test was used to compare values under different conditions, and the differences were considered significant when p ≤ 0.05. When comparing the differences in correlation coefficients across distributions (Fig. 2d), the correlation coefficients (r) were first transformed to a z-score with a fisher z-transformation, z=0.5 * (ln(1+r)-ln(1-r)). Then a z-test statistic (z_observed_) was computed using the formula (z_1_-z_2_)/√((1/(n_1_-3)+(1/(n_2_-3)), where z_1_ and z_2_ are the z-scores of the two distributions, and n_1_ and n_2_ are the corresponding sample sizes. The two distributions were considered significantly different when z_observed_ was greater than 1.96, the critical value for a significance level of 0.05. For the imaging experiments, we used Wilcoxon signed-rank test to evaluate the effects of ambenonium. To evaluate the uniformity in the angular distribution, we used Hodges-Ajne test (Fig. 4g).

## Acknowledgements

We thank Dr. Kevin Briggman for his useful discussions; Tracey Michaels for performing AAV injections and help with mouse colony management; Dr. Ben Murphy-Baum for assistance in writing software routines for image analysis; Zoltan Raics for developing our visual stimulation system, and Bjarke Thomsen and Misugi Yonehara for their technical assistance. Dr. Marla Feller for nGFP mice. Dr. Jamie Boyd for his help with IGOR software for 2P imaging. This work was supported by grants awarded to ‘AM’ (VELUX FONDEN Postdoctoral Ophthalmology Research Fellowship:27786); ‘JMM’ (NIH GM136430 and GM103801); ‘YLL’ (General Research Program of National Natural Science Foundation of China (project 31671118), the NIH BRAIN Initiative (grant U01NS103558), the Beijing Brian Initiative of Beijing Municipal Science & Technology Commission (Z181100001518004), the Junior Thousand Talent Program of China, and by grants from the Peking-Tsinghua Center for Life Sciences and the State Key Laboratory of Membrane Biology at Peking University School of Life Science); ‘DB’ (RO1 EY012793-19); ‘KY’ (Lundbeck Foundation: DANDRITE-R248-2016-2518; R252-2017-1060, Novo Nordisk Foundation, NNF15OC0017252, Carlsberg Foundation, CF17-0085, and European Research Council Starting, 638730) and ‘GBA’ (CIHR 159444).

## Figures and Legends

**Extended Data Fig. 1:**
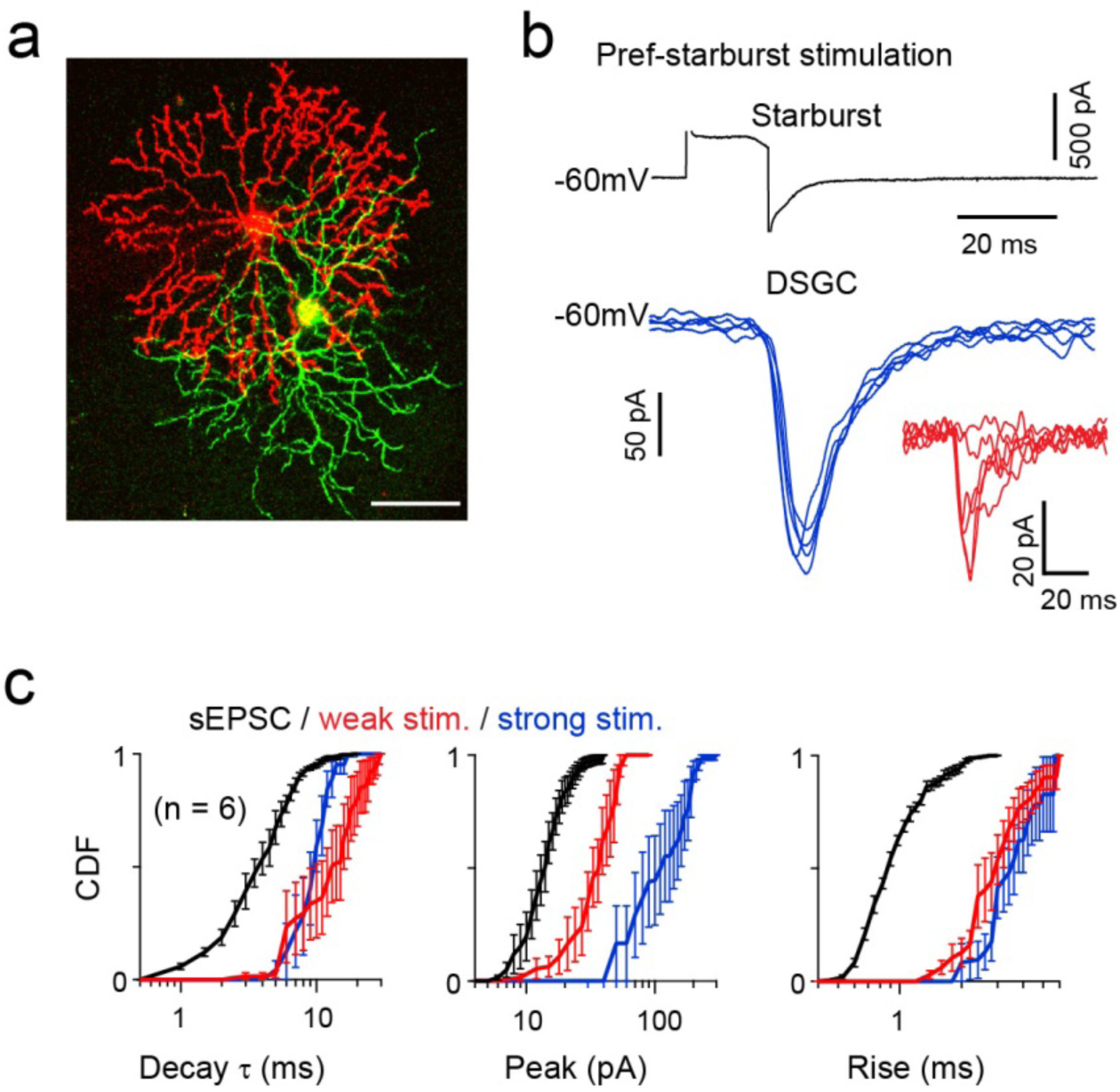
Evoked cholinergic currents are slow to rise relative to spontaneous EPSCs, regardless of the level of stimulation making it difficult to ascertain whether they are mediated by paracrine or synaptic mechanisms. **a.** Two-photon image stack showing the morphology of a connected starburst and a DSGC, recovered after a paired whole-cell patch-clamp recording during which they were dialyzed with an indicator. Scale bar=50μm. **b.** Brief voltage pulses delivered to the starburst (0mV, for 17ms) voltage-clamped at -60 mV, evoked robust cholinergic EPSCs in the DSGC. Weaker pulses (−10mV, for 17ms) evoked smaller amplitude responses, less reliably (Inset). **c.** The kinetics of the evoked EPSCs are slow compared to the sEPSCs events. A comparison of the mean peak (*left*), rise times (*middle*) and decay constant (*right*) of the sEPSCs with evoked EPSCs (red: weak stimulation; blue stronger stimulation; n=6). (Extended Data Table 1; peak: p < 0.001; rise: p < 0.005; decay: p < 0.05). Data represented as mean ± SEM.

**Extended Data Fig. 2:**
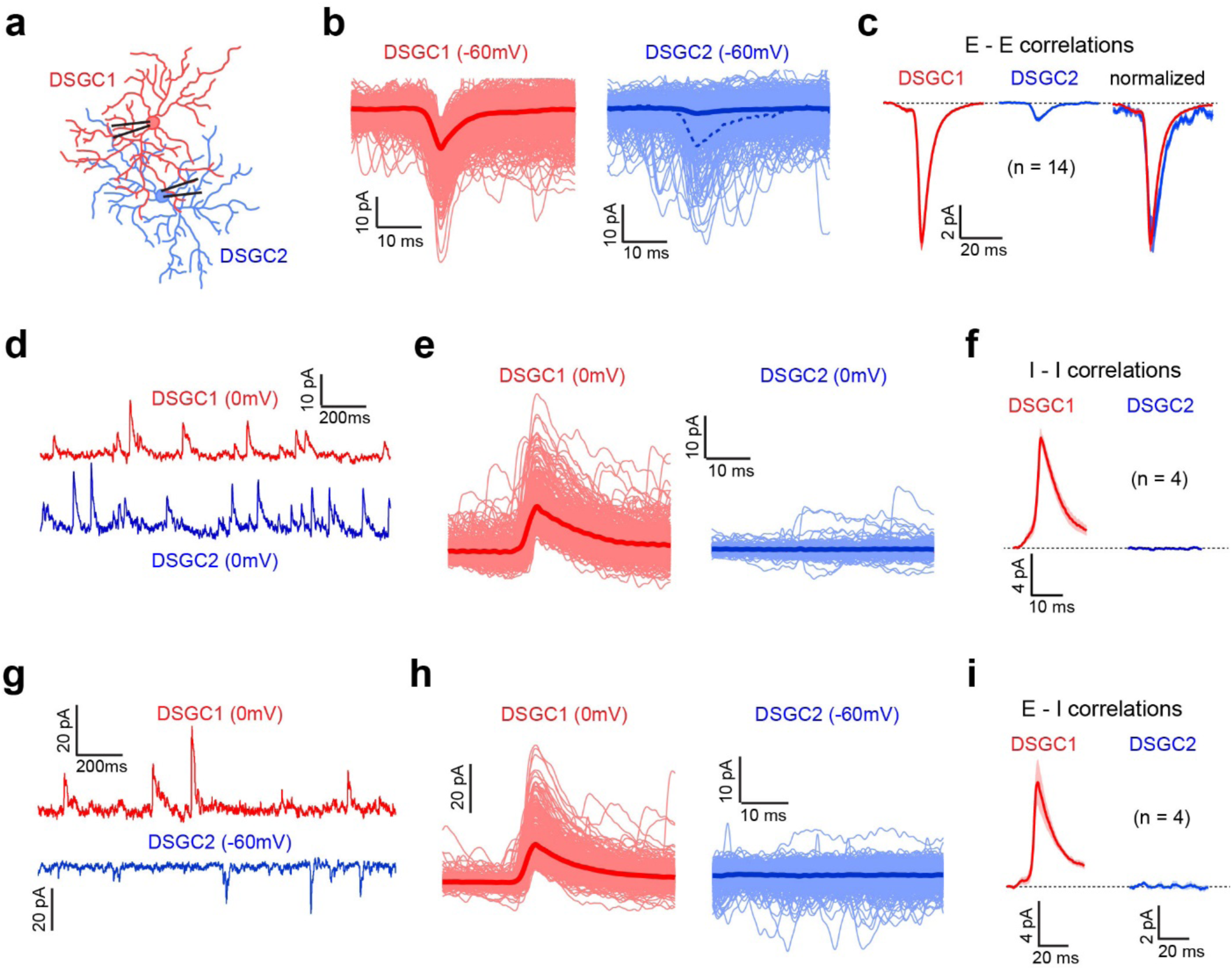
Spontaneous E-E, I-I and E-I correlations in adjacent DSGCs. **a.** Spontaneous EPSCs and or sIPSCs were monitored simultaneously in nearby DSGC with overlapping dendritic fields. **b.** Spontaneous events in a reference DSGC were peak-aligned (red); currents over these same periods as the reference cell, measured in the second DSGC (blue). Dark lines indicate the average current over all the sweeps, and their ratio provides an estimate of the correlation strength (dotted line depicts the average current normalized to the sEPSC in DSGC1). **c.** Spontaneous EPSC and correlated sEPSCs across 7 pairs of DSGCs (for each recording two unique values are obtained by considering each DSGC as a reference cell separately). **d.** GABAergic sIPSCs were simultaneously recorded by voltage clamping both the DSGCs at 0mV. **e.** Aligning currents in DSGC2 to sIPSC in DSGC1 results in an average current that is flat, indicating that they are not correlated. **f.** The average sIPSC (*left*) compared to the correlated GABAergic currents (*right*) in a subset of DSGCs shown in **c** (n=4 pairs). **g-i.** An analysis similar to d-f, but for sIPSCs recorded in one DSGC and sEPSCs in the second DSGC.

**Extended Data Fig. 3:**
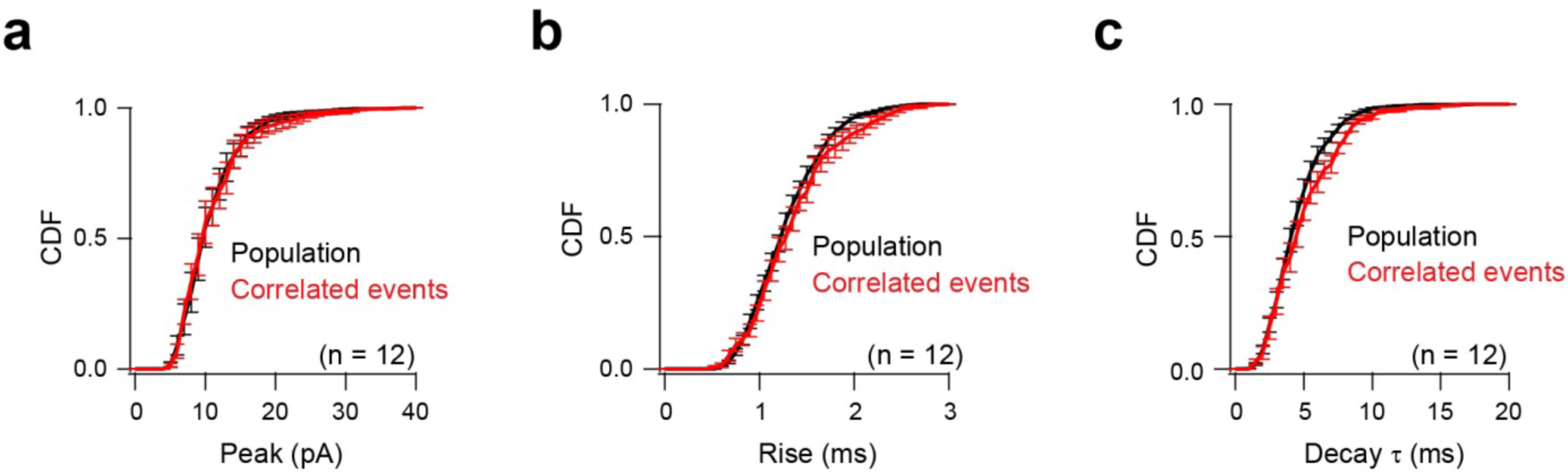
The properties of the sEPSCs that are synchronous in two DSGC are similar to those of the population. **a.** A comparison of the cumulative frequency distributions of the peak of the ACh sEPSC population in DSGCs compared to the sub-population of correlated sEPSCs shown in Fig. 2d. Data represented as mean ± SEM. **b, c.** Similar to a, but for rise times (b) and decay *τ* (c). Data represented as mean ± SEM.

**Extended Data Fig. 4:**
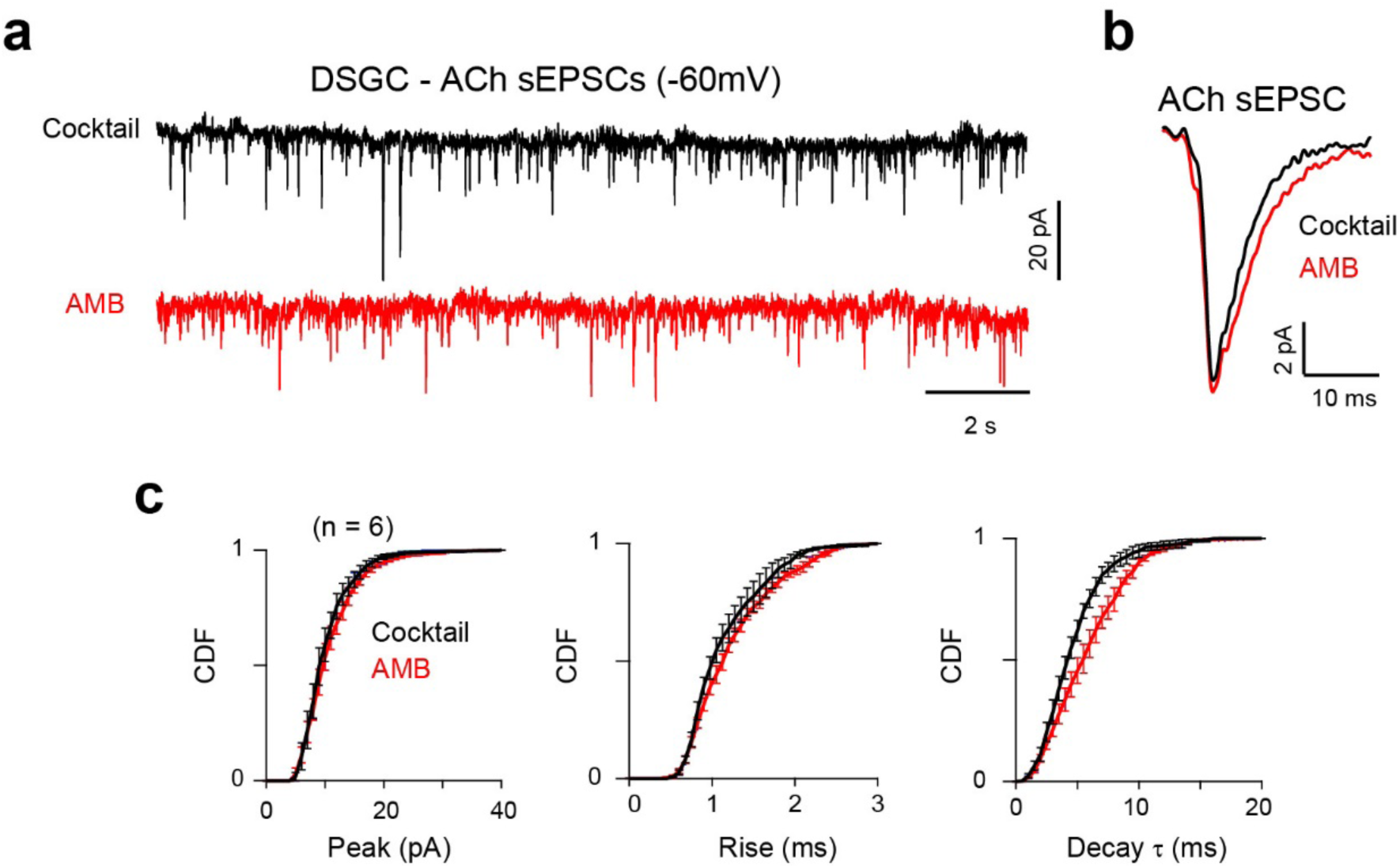
ACh degradation shapes the decay kinetics of cholinergic sEPSC. **a.** Cholinergic sEPSCs measured in a voltage-clamped DSGC, in control (black) and in the added presence of ACh esterase blocker (50 nM ambenonium; AMB; red). **b.** Average sEPSCs waveforms with and without AMB (average obtained from 250, 400 events). **c.** A comparison of the cumulative frequency distributions of the peak (*left*), rise times (*middle*) and decay constants (*τ*) (*right*) of the sEPSCs shown in a and b (n=6 cells). The decay is significantly prolonged in AMB (p < 0.001, paired t-test). Data represented as mean ± SEM.

**Extended Data Fig. 5:**
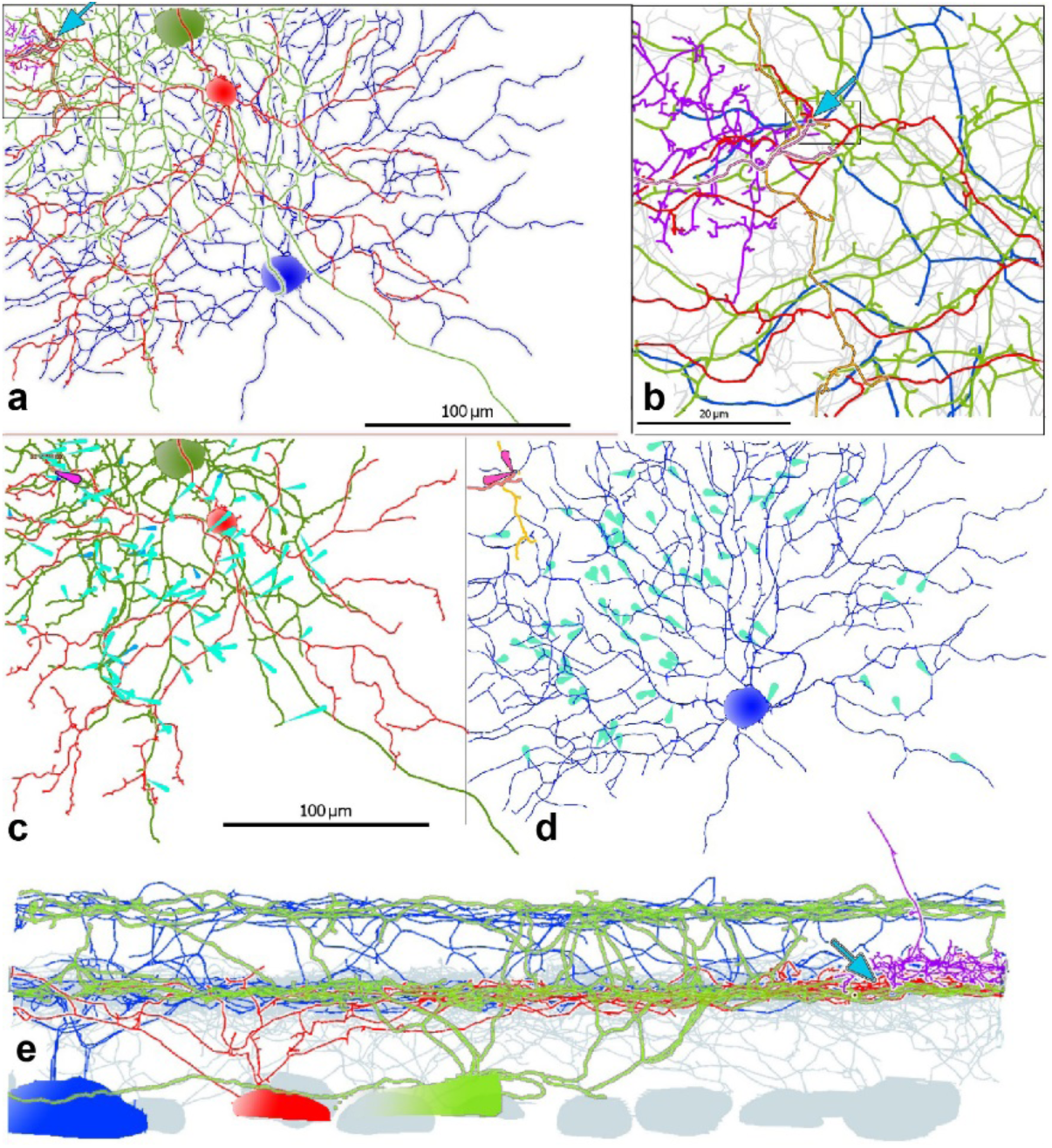
SBEM reconstruction of synapses and contacts between an ON SAC and two postsynaptic ON-OFF DSGCs with opposite preferred directions. **a.** *En face* view of the reconstructed cells. Large blue and green cells are ON-OFF DSGCs. The red cell is an ON SAC. Two other ON SAC dendritic fragments (gold and peach) are also seen entering the volume at upper left. Blue arrow points to the location of the synapse between the red SAC and green DSGC (seen in Fig. 4d). Also shown are the dendrites of a Type 5o bipolar cell (pink), which makes a ribbon synapse with the red SAC and blue DSGC. **b.** Enlargement of the boxed area in a. Blue arrow marks the same synapse as in a. **c, d.** Opposing patterns of asymmetry in SAC contacts onto the two DSGCs. Light blue arrows mark the centrifugal orientation of SAC boutons making synaptic contacts onto these two ganglion cells. The green DSGC receives SAC synapses predominantly from processes coursing away from their parent cell body in a leftward and slight upward direction (c). This is true for the synapse from the red SAC as well, whose orientation marker is highlighted in pink. This cell thus presumably prefers motion down and right in the volume, as illustrated. By contrast, the blue DSGC receives SAC synapses coursing in the opposite direction (d), down and right, and thus would be expected to prefer motion up and left in the volume. The orientation of two SAC processes synapsing onto this cell (peach and gold) is consistent with this, as shown by the pink orientation markers. **e.** Rotated view of all of the same cells, overlaid on the profiles of all reconstructed ON SACs (light grey). The DSGCs show the expected bistratified arbor. The lower (inner) of these two corresponds to the ON SAC plexus. Blue arrow shows the locus of the same synapse.

**Extended Data Fig. 6:**
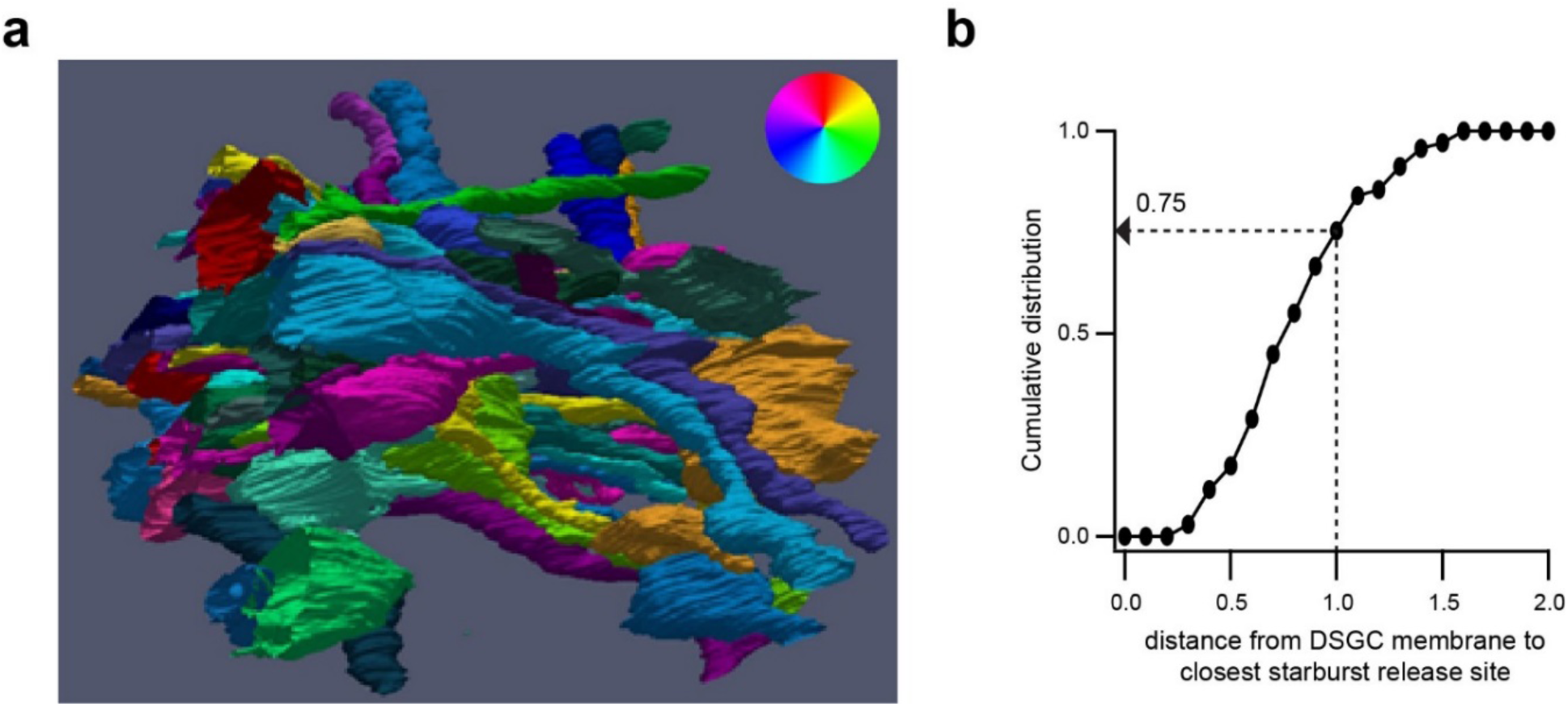
Estimating the average distance of DSGC dendrites from starburst release sites. **a.** An exhaustive reconstruction of the SAC-DSGC network in a small volume (7 × 7 × 7 *µ*m) centered on a dense aggregation of ON SAC processes (same region as Fig. 1a). Among our library, 43 starbursts, 5 ON-OFF DSGCs and 2 ON DSGCs had dendrites that traversed this volume. The orientation (and directional preference) of the starburst dendrites are shown in color. Of the 81 starburst synapses in this volume, roughly a third were made onto ON-OFF DSGCs (n=30; 37%) and another third onto ON SACs (n=28; 34%). The remainder were made onto other RGCs (n=12; 15%) or onto medium or wide-field amacrine cells (n=11; 14%). **b.** A plot showing the fraction of starburst varicosities as a function of distance from the DSGC dendrites in this volume. Note that only starburst varicosities not synapsing onto the examined DSGC were considered for this analysis.

**Extended Data Fig. 7:**
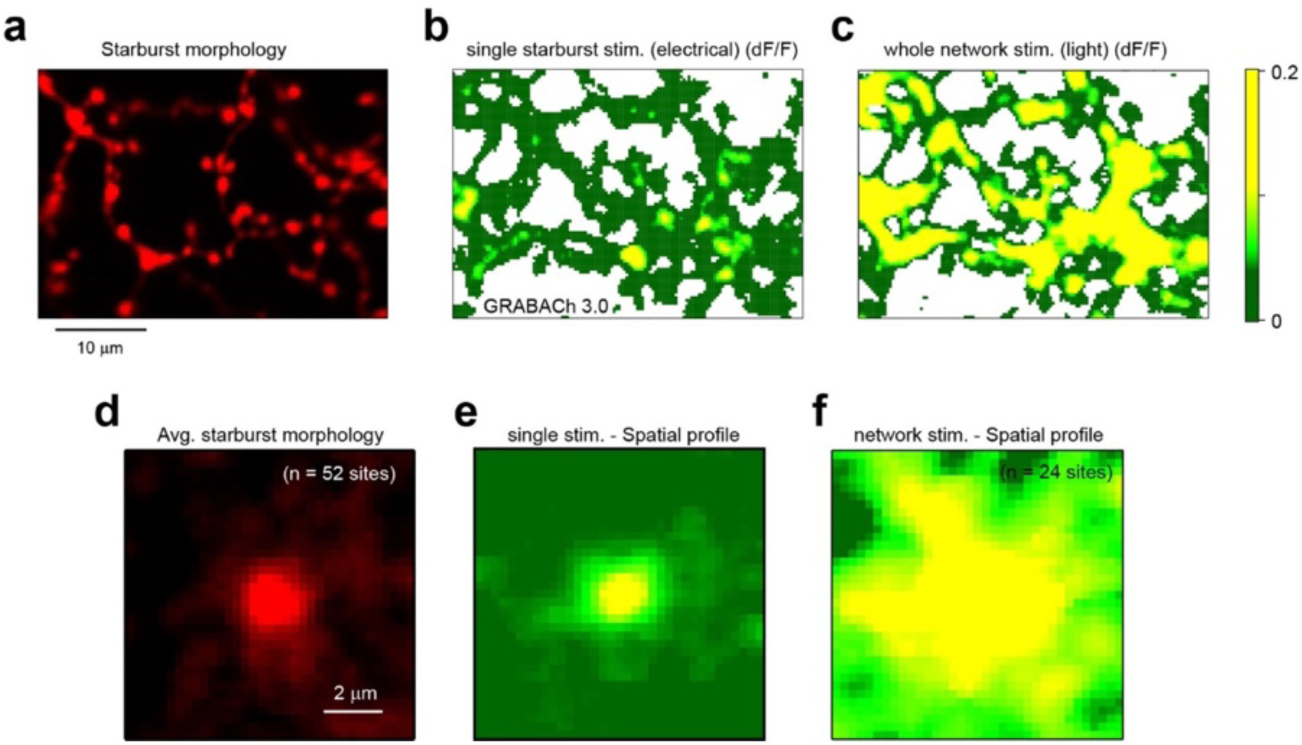
The spatial profile of ACh signals is not limited by the ACh3.0 sensor expression. **a.** An image stack showing the morphology of a starburst (red channel) that was loaded with a red dye through the patch electrode used to stimulate it. **b.** An image of the changes in fluorescence after brief depolarisations of the starburst shown in a (average of 7 trials). The white space indicates regions without ACh3.0 expression. **c.** Changes in fluorescence (average of 3 trials) across the same field of view, evoked by a moving spot appear more widespread. **d.** The average fluorescence in the red channel weighted according by peak ACh3.0 signals evoked by stimulating single starburst (n=52 sites from 8 cells). **e.** The average spatial profile of ACh3.0 signals evoked by single starburst stimulation (same as Fig. 4c). **f.** The average ACh3.0 signal in response evoked by a moving spot (n=24 regions).

**Extended Data Fig. 8:**
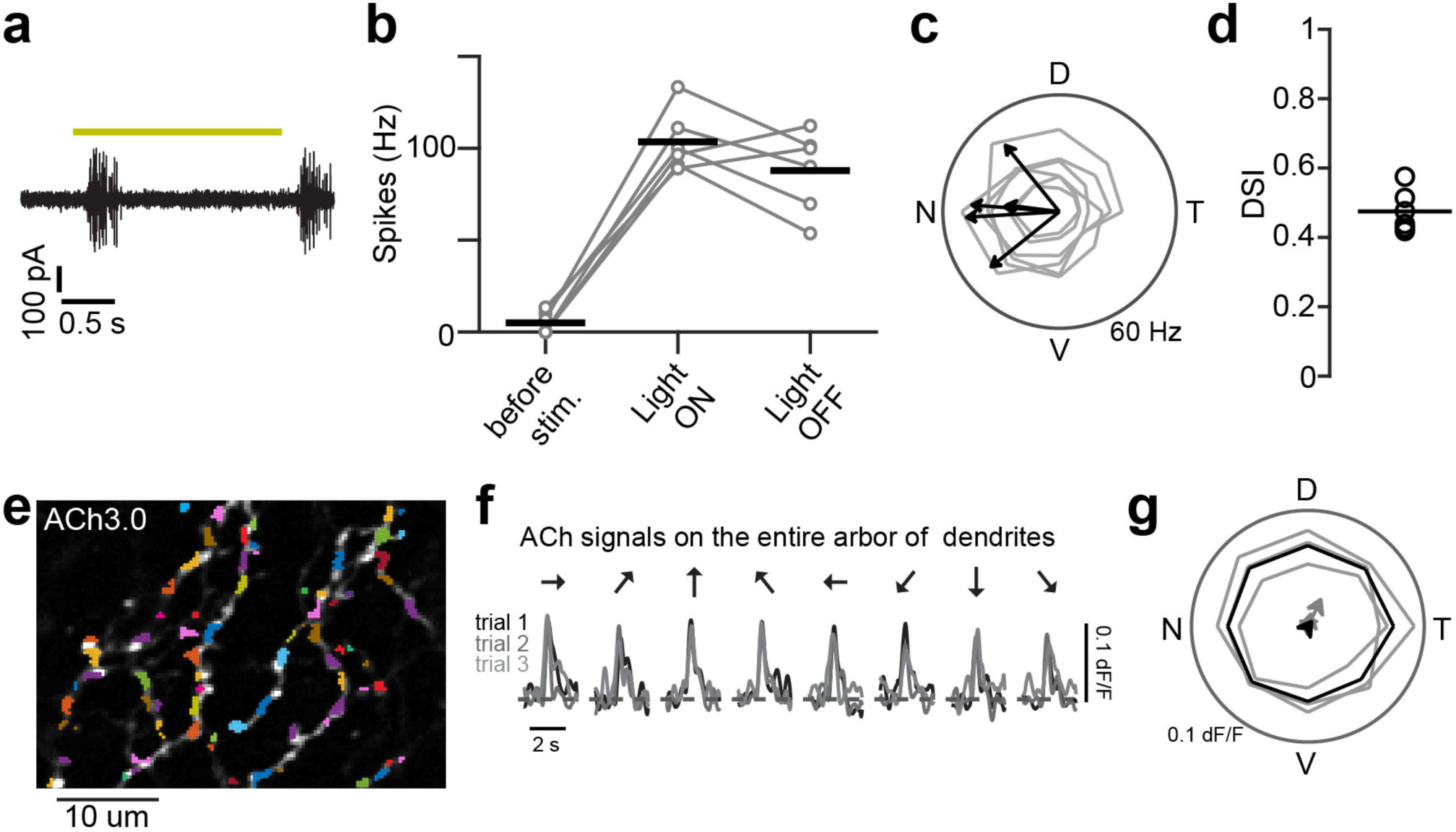
Two-photon acetylcholine imaging with Ach3.0 expressed in genetically labeled nasally-tuned ON-OFF DSGCs. **a.** Example spiking responses to the onset and offset of a static flash stimulus (yellow line indicates 2 s duration of a 300 *µ*m spot, presented at 50 % contrast) recorded from a DSGC labelled with an ACh3.0 sensor in Oxtr-T2A-Cre mouse line. **b.** The peak spiking rates evoked at light ON and OFF (gray, 6 cells; black, average). **c, d.** Directional tuning of the labeled DSGCs (c; gray, 6 cells; arrow, vector sums in the individual cells) in responses to motion stimulus (500 *µ*m in diameter, 1000 *µ*m/s, 50 % contrast). The labeled cells showed directionally-tuned firing with an overall preference for nasal direction (d; circle, 6 cells; black, average; DSI, 0.47 ± 0.03). (D: dorsal; V: ventral; N: nasal; T: temporal). **e.** An example field-of-view (FOV) illustrating the dendritic expression of ACh3.0. The pixels with noise correlation higher than 2.5 during were used to define regions of interests (ROIs; depicted by the different colors) (See Fig. 4e). The detected ROIs with response index higher than 0.5 were classified as ‘responsive’ (see Methods). **f.** Stable ACh responses measured across the entire dendritic arbor of the FOV shown in e (3 sets of 8 trials). These dF/Fs were calculated only from responsive ROIs in the FOV (76 ROIs in e). **g**. The average response measured in **f** does not show a bias towards the preferred direction (nasal direction).

**Extended Data Table 1:** Comparison of spontaneous and evoked response kinetics under different conditions. The table shows the peak, rise and decay time constant of spontaneous and asynchronous events under different conditions. Data are represented as mean ± SEM.

|  | Condition | Peak (pA) | Rise times (ms) | Decay constant (ms) |
| --- | --- | --- | --- | --- |
| Fig. 2k, l | Pref. sIPSC (n = 7) | 12 ± 1 | 1.0 ± 0.1 | 5.4 ± 0.8 |
|  | Pref. aIPSC (n = 7) | 12 ± 1 | 1.0 ± 0.1 | 5.6 ± 0.4 |
|  | Null sIPSC (n = 7) | 11 ± 1 | 1.1 ± 0.1 | 5.1 ± 0.4 |
|  | Null aIPSC (n = 7) | 11 ± 1 | 1.1 ± 0.1 | 4.5 ± 0.6 |
| Extended<br>Data Fig. 1 | sEPSC (n = 6) | 16 ± 2 | 0.99 ± 0.03 | 4.4 ± 0.3 |
|  | Weak stim. (n = 6) | 38 ± 5 | 3.3 ± 0.5 | 15.9 ± 3.8 |
|  | Strong stim. (n = 6) | 131 ± 25 | 7.2 ± 1.3 | 9.8 ± 1.2 |
| Extended<br>Data Fig. 3 | sEPSC whole Population (n = 12) | 11 ± 1 | 1.3 ± 0.1 | 4.5 ± 0.2 |
|  | Correlated sEPSCs only (n = 12) | 11 ± 1 | 1.4 ± 0.1 | 5.0 ± 0.3 |
| Extended<br>Data Fig. 4 | Control sEPSC (n = 6) | 11 ± 1 | 1.2 ± 0.1 | 4.7 ± 0.4 |
|  | AMB sEPSC (n = 6) | 12 ± 1 | 1.3 ± 0.1 | 5.9 ± 0.3 |

